# Signatures of human European Paleolithic expansion shown by resequencing of non-recombining X-chromosome segments

**DOI:** 10.1101/067835

**Authors:** Pierpaolo Maisano Delser, Rita Neumann, Stéphane Ballereau, Pille Hallast, Chiara Batini, Daniel Zadik, Mark A. Joblingt

**Author notes:** Current address: Department of Zoology, University of Cambridge, Cambridge, CB2 3EJ, UK. Current address: Cancer Research UK Cambridge Institute, University of Cambridge, UK. Current address: Institute of Molecular and Cell Biology, University of Tartu, Tartu, Estonia (Pille Hallast). Corresponding author: Prof Mark A. Jobling, Department of Genetics, University of Leicester, University Road, Leicester LE1 7RH, UK. Conflict of Interest:* the authors declare no conflict of interest.

## Abstract

Human genetic diversity in Europe has been extensively studied using uniparentally-inherited sequences (mitochondrial DNA [mtDNA] and the Y chromosome), which reveal very different patterns indicating sex-specific demographic histories. The X chromosome, haploid in males and inherited twice as often from mothers as from fathers, could provide insights into past female behaviours, but has not been extensively investigated. Here, we use HapMap SNP data to identify segments of the X chromosome in which recombination is historically absent and mutations are likely to be the only source of genetic variation, referring to these as Phylogeographically informative Haplotypes on Autosomes and X chromosome (PHAXs). Three such sequences spanning a total of ~49 kb were resequenced in 240 males from Europe, the Middle East and Africa at an average coverage of 181 ×. PHAXs were confirmed to be essentially non-recombining across European samples. All three loci show highly homogeneous patterns across Europe and are highly differentiated from the African sample. Star-like structures of European-specific haplotypes in median-joining networks indicate past population expansions. Bayesian skyline plots and time-to-most-recent-common-ancestor estimates suggest expansions pre-dating the Neolithic transition, a finding that is more compatible with data on mtDNA than the Y chromosome, and with the female bias of X-chromosomal inheritance. This study demonstrates the potential of the use of X-chromosomal haplotype blocks, and the utility of the accurate ascertainment of rare variants for inferring human demographic history.

## Introduction

Studies of the origins and histories of European human populations have been transformed by the availability of next-generation sequencing (NGS), which has given access to the genomes of ancient humans and allowed the unbiased ascertainment of sequence variants in modern populations. Autosomal sequence data from ancient remains have demonstrated discontinuity between Paleolithic hunter-gatherers and Neolithic farmers^1-4^, and more recently have pointed to a later shift due to mass migration from the Pontic-Caspian steppe during the Bronze Age^5-8^. In modern populations, NGS-based studies of uniparentally-inherited markers (the male-specific region of the Y chromosome [MSY] and mitochondrial DNA [mtDNA])^9,10^ have also suggested the importance of recent changes, with marked differences between the two systems being attributed to male-specific Bronze-Age expansion^11^. By contrast, information emerging from resequencing the autosomal genomes of modern individuals has been of limited utility in understanding the events of European prehistory.

The properties allowing MSY and mtDNA to provide useful insights into the past are their haploidy and lack of recombination, which permit demographic reconstruction from haplotypes, and their uniparental inheritance, which provides a sex-specific aspect to the demographic inferences. In this light, the X chromosome represents a potentially useful additional source of information^12^ that has yet to be fully exploited. It is inherited twice as frequently from mothers as from fathers and therefore contains a record biased towards past female behaviours. In males it is haploid, so sequencing in male individuals provides unambiguous phasing of haplotypes, including those bearing rare variants. Finally, it shows high average levels of linkage disequilibrium because most of its length is exempt from crossover in male meiosis. It should therefore be possible to identify segments of the X chromosome that have histories of little or no recombination, to determine their sequences unambiguously in modern male samples using NGS methods, and to use demographic reconstruction to infer female-biased histories.

A number of resequencing studies of X-chromosomal loci have been carried out previously. These have generally surveyed segments of 1-10 kb in global samples^13-21^, and have demonstrated excess gene diversity in African compared to non-African populations, consistent with observations from the autosomes^22^. Some of these studies have chosen segments in which recombination is expected to be low^18-20^, allowing the relatively simple construction of gene trees, while others have not used low recombination rate as a selection criterion, and have yielded haplotypes presenting evidence for multiple recombination events^16^. To the authors’ knowledge, a population-based study of X-chromosomal sequence diversity in Europe has not yet been undertaken.

Here, we select three segments of the X chromosome that show no historical recombination in HapMap data, and use a population resequencing approach to analyse these in multiple European population samples, as well as single Middle Eastern and African population samples. Our findings show that the non-recombining nature of these segments persist, the histories of these X haplotypes in Europe are dominated by Paleolithic expansions, and suggest that larger-scale investigation of the X chromosome will provide further useful insights into the European past.

## Materials and Methods

### Samples

Two hundred and forty DNA samples were analysed, comprising twenty randomly chosen males from each of twelve populations. The list of samples including population origins is reported in Table S1 and additional details were previously described^11^. An additional thirteen unrelated male samples (see Supplementary Material) from six populations were analysed from the Complete Genomics sequence dataset^23^.

### PHAX identification process

PHAXs were originally defined using publicly available SNP data from the HapMap project^24^, release 21, for four population samples: the Centre d’Etude du Polymorphisme Humain (CEPH) collection in Utah, USA, with ancestry from Northern and Western Europe (CEU), the Yoruba from Ibadan, Nigeria (YRI), Han Chinese in Beijing, China (CHB) and Japanese in Tokyo, Japan (JPT). Haplotypes were inferred using PHASE^25^ and measures of LD were downloaded from the HapMap web site, release 16c.

Historically non-recombining regions were identified as non-overlapping series of at least three adjacent SNPs where each pair had a |D’| value of 1 in each of the three samples CEU, YRI and JPT+CHB, and for which only three of the four possible 2-allele haplotypes were observed in the entire sample set, including the ancestral haplotypes (inferred from chimpanzee data), whether or not these were themselves observed. See Supplementary Material for additional details.

### Amplicon sequencing; data analysis, variant calling and filtering

~10 ng of genomic DNA was used for amplification of the three chosen PHAXs via ten amplicons, through Polymerase Chain Reaction (PCR). Amplicons were electrophoresed and quantified on 0.8% (w/v) agarose gels, and pooled at equimolar concentrations per sample. For each sample, 100 ng of amplified pooled DNA was used for library preparation, which was performed with the Ion Xpress™ Plus gDNA Fragment Library Preparation kit (Life Technologies). Size-selection was done using Agencourt AMPure XP beads (Beckman Coulter). Ion Xpress™ barcodes were used to tag individual libraries, which were quantified using the 2100 Bioanalyzer (Agilent Technologies) and the Agilent High Sensitivity DNA Kit. An equimolar pool of libraries was prepared at the highest possible concentration, then fractionated on a 2% (w/v) NuSieve 3:1 agarose gel. Final size-selection and clean-up were performed with the Zymoclean™ Gel DNA Recovery Kit (Zymo Research). Template preparation was carried out using the Ion Xpress™ Template 200 Kit (Life Technologies), prior to sequencing on an Ion Torrent PGM platform with 200-bp reads using the Ion PGM™ Sequencing 200 Kit (Life Technologies) and the Ion 316 chip v1.

Reads were mapped to the human reference sequence (hg19) using TMAP software implemented in the Ion Alignment plugin 3.2.1 (Torrent Suite™ Software 3.2.1). Local realignment and duplicate read marking were carried out with the Genome Analysis Tool Kit (GATK)^26^ and picard v1.94 (http://picard.sourceforge.net/) respectively. All sites were called using SAMtools 0.1.19^27^ and filtering was done with in-house scripts. A total of 49,070 bp were called, including 419 raw variants from 240 samples. Following filtering, 297 variants and 238 samples were retained.

***In silico*** validation was done using Complete Genomics whole-genome sequence data (http://www.completegenomics.com/public-data/69-genomes/: 9 samples) and Illumina sequence-capture data^28^ using only shared called sites in the PHAXs sequenced here (220 samples). Based on the Complete Genomics comparison, the false-positive rate was 0.0005% and false-negative rate zero, while via the Illumina comparison the false-positive rate was zero and the false-negative rate 0.00003%. Further details on filtering and data analysis are reported in the Supplementary Material.

### Intra-and inter-population diversity

Haplotype diversity^29^, Tajima’s D^30^, and Fu’s Fs^31^ were calculated for each PHAX per population using Arlequin v3.5^32^.

Genetic differentiation between populations was measured with the molecular index ϕst^33^, computed with Arlequin v3.5^32^.

### Networks and Bayesian Skyline Plots

Relationships between different haplotypes were displayed in median-joining networks^34^, implemented in Network 4.6 (http://www.fluxus-engineering.com/sharenet.htm). Ancestral state for each site was defined by comparison with the chimpanzee reference sequence.

Bayesian Skyline Plot (BSP) analyses were performed using BEAST v 1.8.0^35^. Markov chain Monte Carlo (MCMC) samples were based on 200,000,000 generations, logging every 10,000 steps, with the first 20,000,000 generations discarded as burn-in. Traces were manually evaluated and inspected using Tracer v 1.6 (http://beast.bio.ed.ac.uk/software/tracer/). A piecewise linear skyline model with ten groups was used with a Hasegawa, Kishino and Yano (HKY) substitution model^36^ and a strict clock with a mean substitution rate of 6.59 × 10^-10^ mutations/nucleotide/year (details of mutation rate estimation are provided in Supplementary Material). A generation time of 30.8 years was used^37^.

### TMRCA estimation

TMRCA estimation for specific haplotype clusters was performed using the rho statistic^38,39^, using Network 4.6 (http://www.fluxusengineering.com/sharenet.htm). This analysis was performed on individual PHAXs with a scaled mutation rate of 41,171, 304,525 and 263,309 years per mutation for PHAX 5574, 3115 and 8913 respectively. These estimates were based on the mutation rate (6.59 × 10^-10^ mutations/nucleotide/year) and the number of nucleotides for each PHAX (36,857, 4983 and 5763 for PHAX 5574, 3115 and 8913 respectively).

## Results

### Selection of non-recombining X-chromosomal regions

We used genomewide HapMap data and linkage disequilibrium (LD) analysis (Supplementary Material) to identify regions showing no evidence of historical recombination in 210 unrelated individuals from the four populations CEU (Utah Residents with Northern and Western European ancestry), YRI (Yoruba in Ibadan, Nigeria), JPT (Japanese in Tokyo, Japan) and CHB (Han Chinese in Beijing, China). Regions passing our LD filters were designated Phylogeographically Informative Haplotypes on the Autosomes or X chromosome (PHAXs).

For the purposes of this study, we applied additional filters to the set of PHAXs identified on the X chromosome in order to obtain a set of informative, independent and putatively neutrally-evolving markers. Candidate regions for resequencing were chosen to be: (i) free of genes; (ii) separated from the nearest known or predicted gene by at least one recombination hotspot; (iii) lacking in segmental duplications, for ease of sequence interpretation; and (iv) possessing an ortholog in the chimpanzee genome^40^, for convenience of ancestral state determination. PHAXs passing these filters were sorted by the number of haplotypes defined by SNPs in the CEU sample^24^, given that the focus of our study was on European populations. The top three regions, spanning ~49 kb in total (Table 1, Figure 1, Supplementary Material), were then divided into ten ~5-kb amplicons (including short overlaps) for resequencing.

**Table 1:** Features of PHAXs analysed.

| PHAX | Coordinates<br>(hg19) | Length<br>(bp) | No. of<br>haplotypes<br>defined by<br>SNPs<sup>a</sup> | No. of<br>HapMap<br>(PII) SNPs<br>(CEU) | No. of non-HapMap<br>SNPs<sup>b</sup> |
| --- | --- | --- | --- | --- | --- |
| 3115 | chrX:<br>39,519,963-39,524,945 | 4983 | 5 | 6 | 2 |
| 5574 | chrX:<br>94,544,032-94,582,132 | 38,101 | 7 | 10 | 54 |
| 8913 | chrX:<br>141,656,627-141,662,612 | 5986 | 5 | 5 | 44 |

<sup>a</sup> HapMap phase I/II populations (CEU, CHB, JPT and YRI).
<sup>b</sup> From dbSNP 152, insertion/deletions not included.

**Figure 1:**
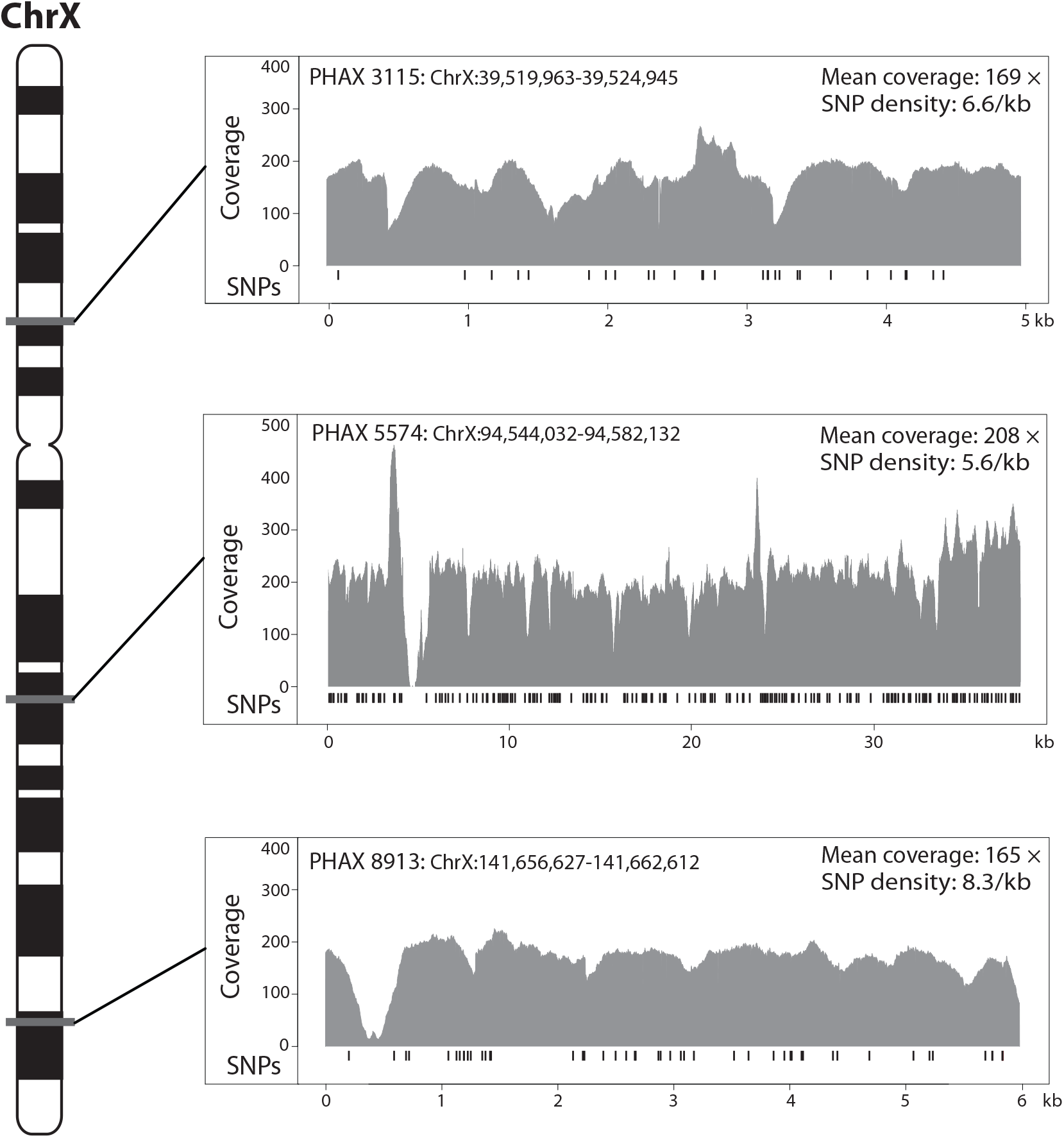
Location, sequence coverage, and SNP distributions for the three PHAXs studied. The approximate positions of the three studied PHAXs are shown on the X-chromosomal ideogram to the left. To the right the three panels show the average coverage of sequence reads across samples per site and the position of SNPs (vertical ticks below the coverage plots) in each PHAX. Mean coverage and SNP density are also shown.

### Genetic diversity and data summary

We used sequencing via the IonTorrent platform to assess the genetic diversity of the three X-chromosomal PHAXs in a total of 240 men from Europe, the Middle East and Africa (Table S1). Mean coverage was 181 × and we called all sites (ignoring indels) having ≥10 × coverage (Figure 1). SNPs were validated *in silico* by comparison with published whole-genome sequences (http://www.completegenomics.com/public-data/69-genomes/), and with a subset of the same samples and target regions sequenced using Illumina technology^28^. The high coverage and high threshold for variant calling led, respectively, to very low false-negative and false-positive rates (Supplementary Material). We ascertained 297 high-quality SNPs in total, which defined 29, 78 and 30 distinct haplotypes respectively for PHAX 3315, 5574 and 8913 (Tables S2-S4). Fifty-eight of the SNPs (19.5%) were not previously reported in dbSNP build 138, and over half (172; 57.9%) were singletons (Figure 2), i.e. unique in the dataset. PHAXs 3115 and 5574 were significantly enriched in singletons (Fu and Li’s D test -5.48 and -7.88 respectively with both p-values <0.02). SNP density varies among the three PHAXs and the difference is marginally significant (Chi Square test with Yates’ correction, Chi Square = 6.024, p = 0.045) suggesting different evolutionary histories for these loci. This was also supported by the significant pairwise differences between the distributions of haplotype diversity of the three PHAXs (Kolmogorov-Smirnov test p-values 0.000084, 0.03 and 0.03 for the comparisons PHAX 5574 vs PHAX 8913, PHAX 5574 vs PHAX 3115 and PHAX 3115 vs PHAX 8913 respectively).

**Figure 2:**
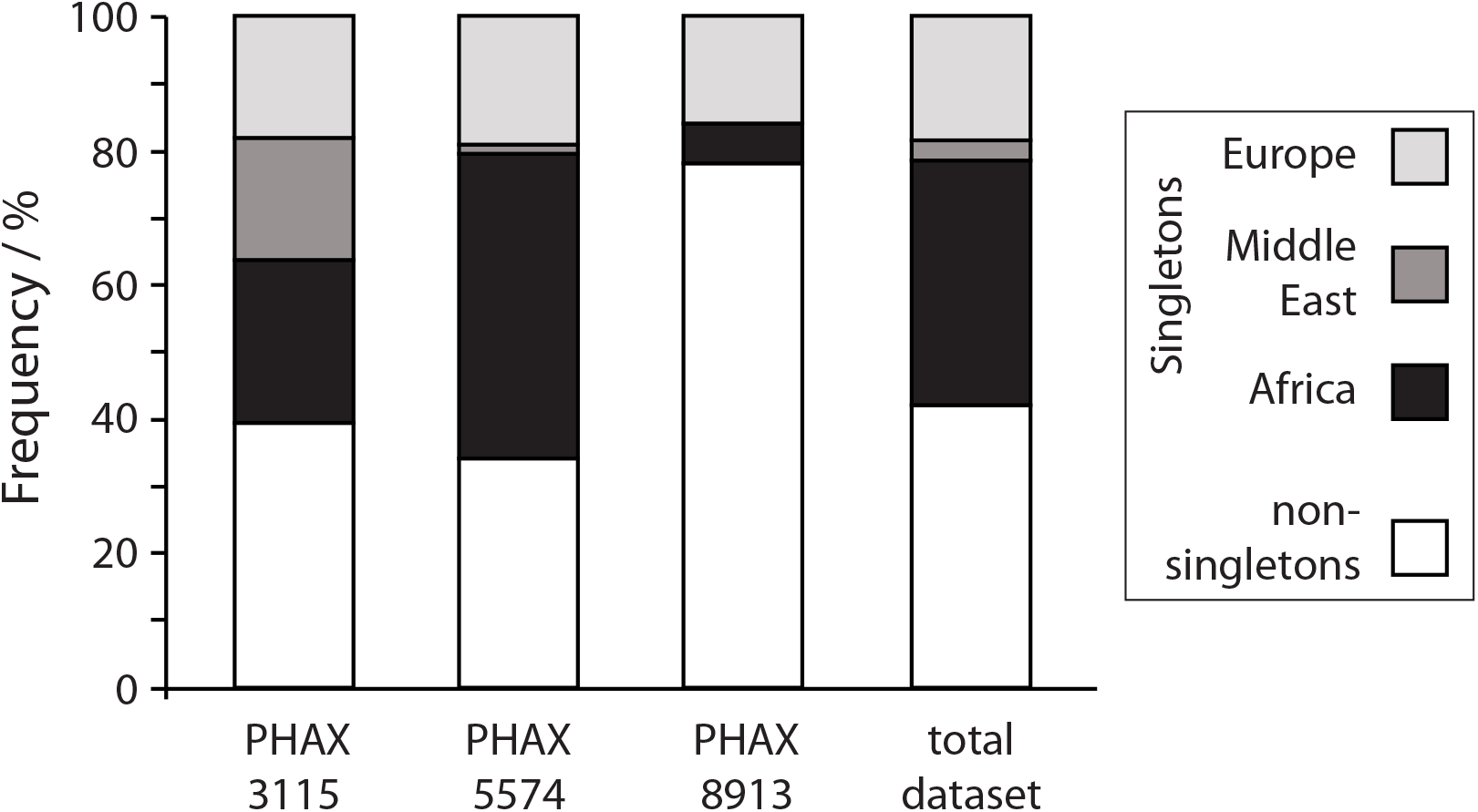
Distribution of singleton and non-singleton variants. Histogram showing distribution of variant types by PHAX, and by meta-population. Africa is represented by YRI (Yoruba from Ibadan, Nigeria); Middle East by Palestinians; Europe by the remaining ten populations.

### Descriptive analyses

Population structure was investigated by performing a Principal Components Analysis (PCA) based on haplotype frequencies within populations. Overall, the first two PCs combined together explain a total of 32% and 61% of variance for the three PHAXs. When all populations were analysed together, the YRI sample was consistently separated from non-African populations in all three cases (Figure S1). To better examine the population structure of non-African populations (Figure S2), the YRI sample was removed in each case, while the Palestinian population was omitted for PHAX 3115 only, since it was an outlier (Figure S2a). All plots show a lack of specific clustering and structure. This pattern was confirmed by the pairwise ϕst matrix (Table S5), which shows strong differentiation between the YRI and other samples, but similar genetic distances among all non-African populations. Overall, genetic distances do not suggest significant population structure within Europe.

### Phylogeographic analysis

Haplotype frequencies were plotted per population according to their geographic locations (Figure 3). For all three PHAXs the YRI sample shows a high number of haplotypes at low and intermediate frequencies (up to a maximum of 18 haplotypes from 20 individuals for PHAX 5574). In Europe, populations carry a few haplotypes at high frequencies and many at low frequencies (0.05). Some local geographical patterns in haplotype distributions can also be observed. For example, haplotype h2 of PHAX 3115 (Figure 3a) is not present among the YRI but is relatively frequent in Europe while haplotype h3 (Figure 3a) is at relatively high frequency in Central and Western Europe, but also persists at low frequency in Middle East and the YRI. Such patterns are more pronounced for PHAX 5574 and PHAX 8913. Haplotype h2 of PHAX 5574 (Figure 3b) is present only in Europe and the Middle East with its highest frequency in Ireland. PHAX 8913 shows a common haplotype (h1) with frequency ranging from 0.6 to 0.8 in Europe that is absent from the YRI (Figure 3c) - features that could indicate a founder effect associated with European colonisation.

**Figure 3:**
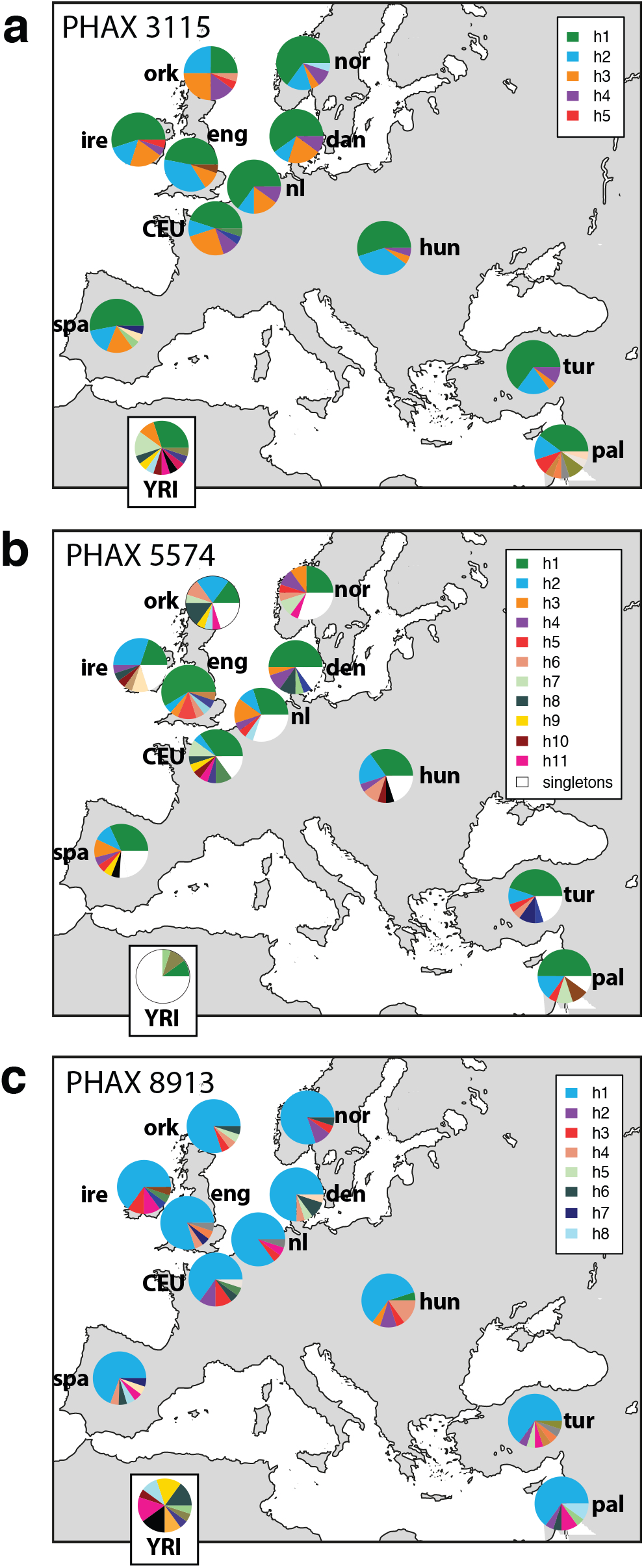
Population distributions of haplotypes. Maps showing distributions of haplotypes for each PHAX, indicated by coloured sectors in pie charts. At the bottom of each panel the distribution for the YRI population is also shown. (a) PHAX 3115: the key indicates non-singleton non-African haplotypes (h1-5). (b) PHAX 5574: the key indicates haplotypes (h1-11) present in three or more non-African individuals; white sectors in pie charts correspond to all singleton haplotypes, which are numerous for this PHAX. (c) PHAX 8913: the key indicates non-singleton non-African haplotypes (h1-8). Population abbreviations are as follows: CEU: Utah residents with Northern and Western European ancestry from the CEPH collection (French); den: Danish; eng: English; nl: Dutch; hun: Hungarian; ire: Irish; nor: Norwegian; ork: Orcadian; pal: Palestinians; spa: Spanish; tur: Turkish; YRI: Yoruba from Ibadan, Nigeria.

To better understand the phylogenetic relationships among haplotypes, and to provide some insight into the reliability of the chosen PHAXs as non-recombining segments, we generated a median-joining network^34^ for each PHAX (Figures 4, S3). Only one reticulation was observed for PHAX 5574 (Figure 4) involving one nucleotide in a private Spanish haplotype, which could be compatible with either a recurrent mutation or a recombination event. This network also displays a pronounced ‘star-like’ structure (Figure 4), in which two major haplotypes show high frequencies with several low-frequency haplotypes linked to them by one or two mutational steps; this seems compatible with a demographic expansion. One of the major haplotypes is h2 (in blue in Figure 3b) while the other haplotype shared across all populations is h1 (in green in Figure 3b). The majority of private YRI haplotypes lie outside the star-like part of the network. Two reticulations exist in the PHAX 3115 network (Figure S3a). These both involve unique YRI haplotypes that interrupt the PHAX, while the segment still remains as an unbroken LD block in non-African populations. The star-like structure is less pronounced for PHAX 3115 compared to the other two PHAXs (Figure 4; Figure S3b), but most branches are one or two mutational steps in length, with one branch carrying four mutation events. PHAX 8913 shows no reticulations in its network, suggesting that this region remains an LD block in the larger dataset, lacking either recombination events or recurrent mutations (Figure S3b). A high-frequency haplotype (h1) shared by all populations in the dataset was found with several haplotypes at lower frequency linked to it only by single mutational events. This structure resembles the PHAX 5574 network and also seems compatible with a demographic expansion generating many haplotypes at low frequencies Overall, the few reticulations found in the three PHAXs, mainly involving African haplotypes, confirm that such sequences remain useful haplotypic markers even when analysed in a larger European dataset compared to the HapMap Phase II data used for their definition. Rare alleles found in non-African populations did not disrupt the LD blocks, and PHAXs still behave according to their original definition. Moreover, the general star-like structure found in all the three networks seems compatible with a demographic expansion that has generated many haplotypes at low frequencies branching from a few common haplotypes that are widely spread across populations.

**Figure 4:**
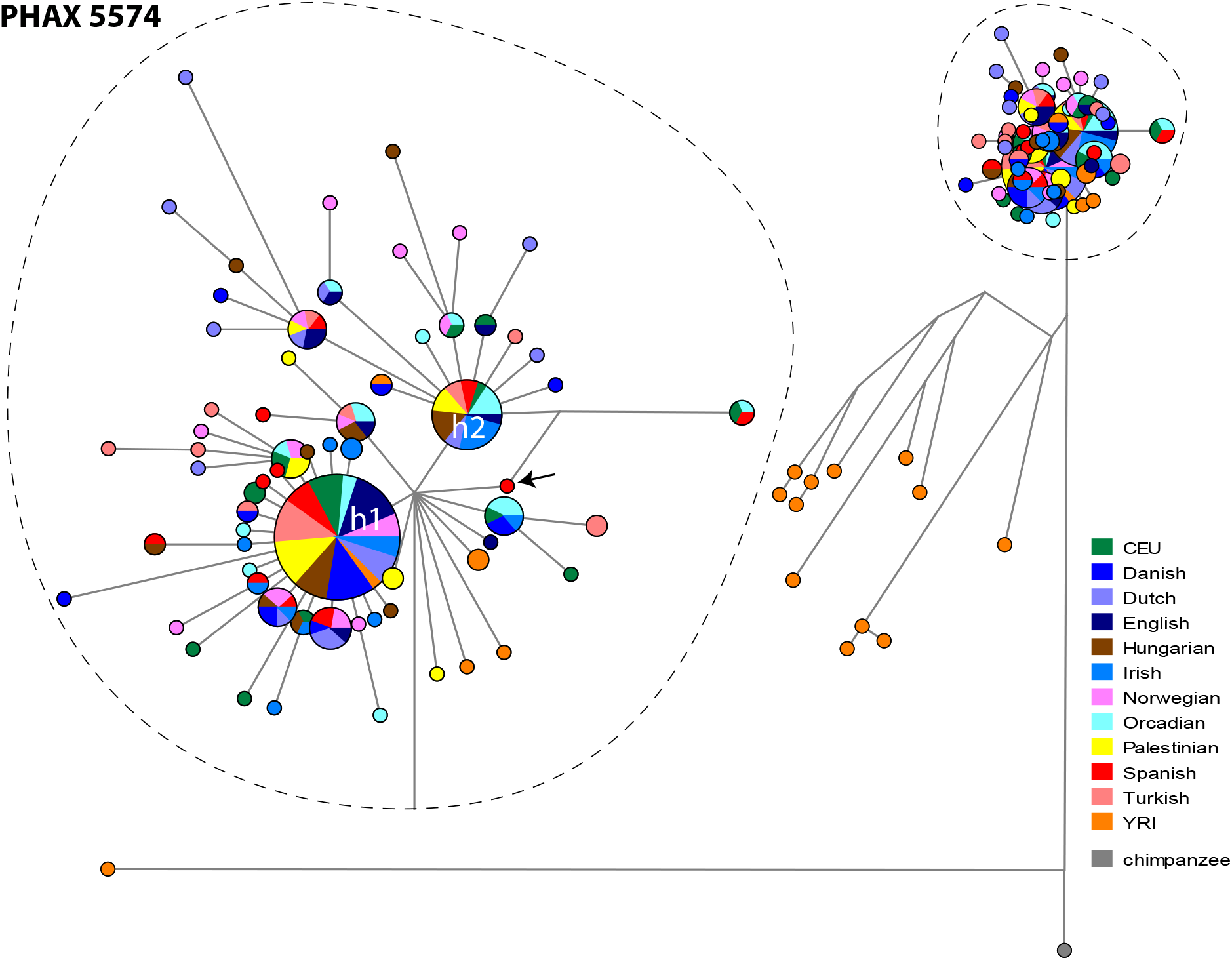
Median-joining network for PHAX 5574 showing population distribution of haplotypes. Circles represent haplotypes with area proportional to frequency, and lines between them represent SNP mutational steps between haplotypes. Populations are indicated by colours as shown in the key to the right. The major haplotype cluster is magnified in the dotted ellipse to the left for clarity, and the private Spanish haplotype involved in a reticulation is highlighted by an arrow. Haplotypes h1 and h2, mentioned in the text, are indicated. Networks for the other two PHAXs can be found in Figure S3.

### Demographic inferences

Bayesian Skyline Plots (BSPs) were produced to analyse the demographic signal in Europe suggested by each PHAX. For this analysis European populations (excluding YRI and Palestinians) were grouped together based on the low genetic distances suggested by the ϕst matrix and the absence of population structure indicated by the PCAs. The BSPs show an increase in effective population size of more than one order of magnitude (Figure 5), consistently across the three PHAXs. This increase starts around 20 KYA (thousand years ago) and becomes more pronounced around 18-15 KYA. PHAX 5574 shows a more constant pattern until 10 KYA, when the increase in effective population size becomes more marked. Overall, the patterns shown by the three PHAXs are compatible with an expansion in Europe starting around 20 KYA. By contrast, the YRI sample (Figure S4) is characterised by flat BSPs that do not suggest any demographic change in effective population size. A similar pattern is seen for the Palestinian population (Figure S5), with BSPs failing to show any strong demographic change. For both the YRI and Palestinian samples, all three PHAXs suggest comparable estimates of modern effective population size.

**Figure 5:**
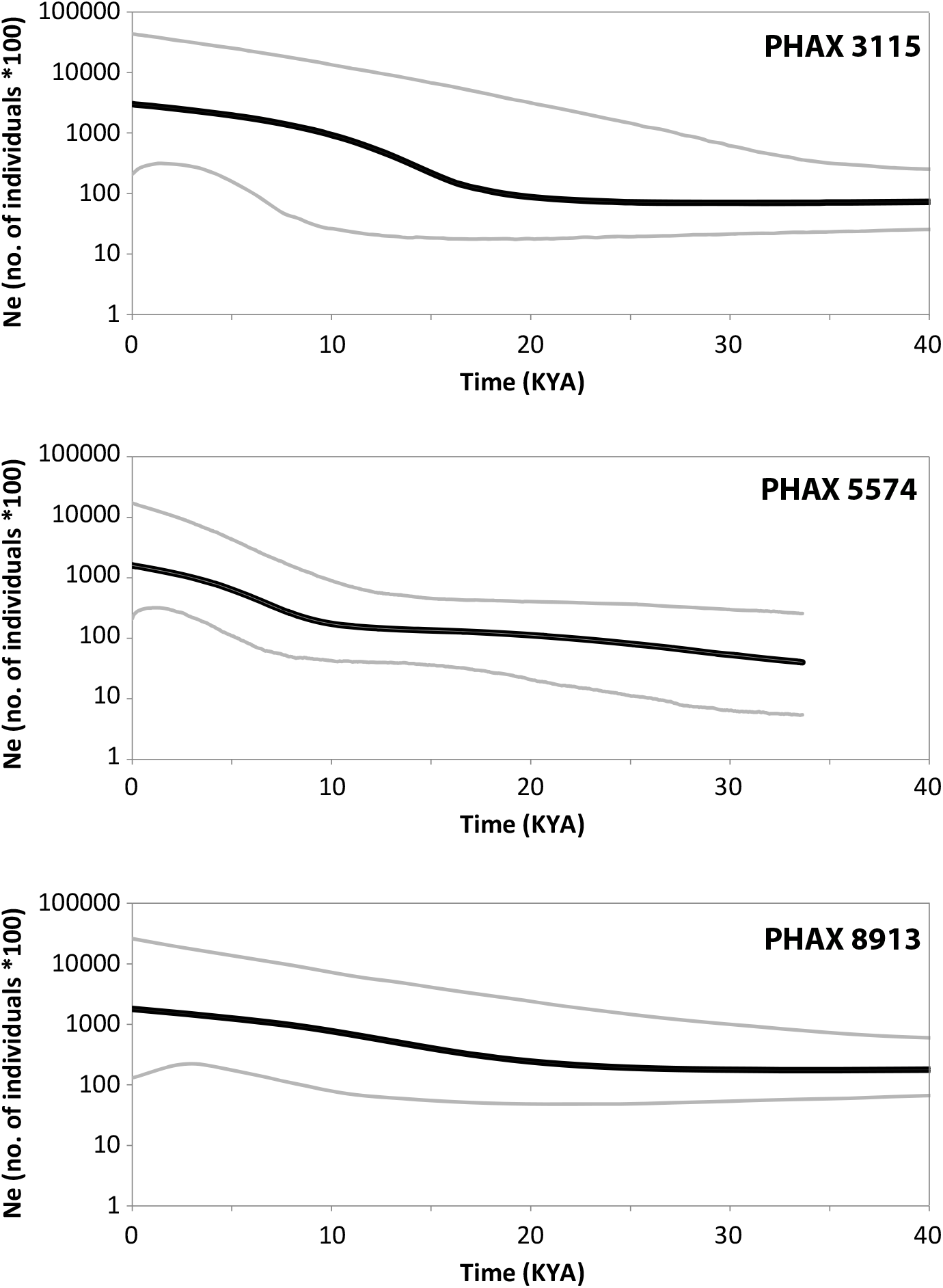
Bayesian Skyline Plots in European populations for the three PHAXs. Thick black lines indicate the median for effective population size (N_e_) and thinner grey lines show 95% higher posterior density intervals. For BSPs based on the YRI and Palestinian population samples, see Figures S4 and S5.

TMRCA (time to most recent common ancestor) was calculated for specific haplotype clusters for each PHAX. Clusters were chosen based on the structures of networks - nodes of specific interest (such as the ancestral node) and prominent ‘star-like’ structures (Figure S6). Mean estimates with standard deviation are reported in Table 2. All the ancestral nodes across the three PHAXs are dated between ~900 KYA and ~1.6 MYA. Cluster 1 in PHAX 3115 was dated ~46.8 KYA; this estimate is in agreement with two other clusters, cluster 2 (PHAX 5574) and cluster 1 (PHAX 8913), which have TMRCA estimates of ~44.85 KYA and ~44.82 KYA respectively. Cluster 3 in PHAX 5574 is the youngest across the dated clusters showing TMRCA ~21.8 KYA, reflecting the very pronounced ‘star-like’ structure in the network. Cluster 1 in PHAX 5574 divides almost all non-African samples from the African ones and has a TMRCA ~68.8 KYA.

**Table 2:** TMRCA estimates.

|  | **TMRCA/YBP** | **s.d./years** |
| --- | --- | --- |
| **PHAX 3115** |  |  |
| Ancestral node | 914,855 | 401,391 |
| Cluster 1 | 46,850 | 14,055 |
| **PHAX 5574** |  |  |
| Ancestral node | 1,194,432 | 212,395 |
| Cluster 1 | 68,801 | 28,059 |
| Cluster 2 | 44,847 | 14,500 |
| Cluster 3 | 21,779 | 5731 |
| **PHAX 8913** |  |  |
| Ancestral node | 1,590,917 | 489,355 |
| Cluster 1 | 44,819 | 18,261 |

For cluster definition, see Figure S6.
TMRCA: time to most recent common ancestor; YBP: years before present; s.d.: standard deviation

## Discussion

Of the ‘odd couple’ of the human sex chromosomes, the Y chromosome has received most attention in population genetics to date, because of its male specificity and the consequences for its mutation processes of its unusual state of constitutional haploidy. But the X chromosome is arguably just as interesting and strange, showing only one copy in males, female-biased mutation processes, and the unique phenomenon of X-inactivation.

The X chromosome also shows reduced crossover activity compared to autosomes, because the non-pseudoautosomal majority (98%) of the chromosome is recombinationally active in only one sex - females. We therefore expect it to contain relatively long segments that show limited historical recombination activity, and can be treated as simple haplotypes in evolutionary studies. The fact that males are haploid for the X chromosome means that studying such segments in males will provide reliably phased haplotypes, even when variants within these are very rare in the population. Here, we have investigated this idea by targeting three candidate X-chromosomal segments based on SNP data that indicate no evidence of historical recombination in the four HapMap Phase I populations. We identified such segments genome-wide, and have designated them Phylogeographically Informative Haplotypes on the Autosomes and X chromosome (PHAXs).

High-coverage resequencing of the three PHAXs here, which cover 49 kb in total, reveals 297 SNPs in a sample of 240 males belonging to 12 populations, 11 of which are European or Middle Eastern. Almost 58% of SNPs are high-quality singletons suggesting that a resequencing approach is crucial for an accurate ascertainment of rare variants which would have been lost with other techniques such as SNP arrays. Low-frequency sites provide vital information to finely assess and reconstruct recent demographic history. Network analysis shows that the absence of recombination generally persists in this independent sample in which variants are well ascertained. Two haplotypes showing evidence for recombination are found among the 20 males in the YRI population, while among 220 European and Middle-Eastern chromosomes, one haplotype shows a single variant that requires recurrent mutation or recombination as an explanation. When additional available high-coverage sequence data from thirteen samples sequenced by Complete Genomics are added to our dataset, this pattern persists. HapMap Phase I data therefore seem a reliable source of information about non-recombining regions for evolutionary studies.

Unsurprisingly, the highest genetic diversity is found in the YRI population sample, consistent with genome-wide data on African vs non-African diversity^22^. This can be seen in summary statistics (Table 3; Figure 2) and in the distribution of haplotypes in the networks (Figure 4; S3). In the non-African samples diversity is lower, and the patterns of haplotypes in network analysis are more star-like, suggesting past population expansions.

**Table 3:** Genetic diversity of the three PHAXs.

|  |  | PHAX3115 | PHAX5574 | PHAX8913 |
| --- | --- | --- | --- | --- |
| Number of segregating sites | Europe | 15 | 64 | 42 |
|  | YRI | 16 | 142 | 35 |
|  | Total | 33 | 214 | 50 |
| Number of singletons | Europe | 6 | 38 | 7 |
|  | YRI | 8 | 97 | 3 |
|  | Total | 20 | 141 | 11 |
| Number of haplotypes | Europe | 13 | 54 | 20 |
|  | YRI | 12 | 18 | 10 |
|  | Total | 29 | 78 | 30 |
| Average $\phi_{st}$ | Europe | 0.017 | 0.010 | 0.018 |
|  | YRI | 0.05 | 0.243 | 0.477 |
|  | Total | 0.023 | 0.048 | 0.096 |
| Nucleotide diversity ($\pi$) | Europe | 0.072 +/- 0.043 | 0.012 +/- 0.007 | 0.054 +/- 0.032 |
|  | YRI | 0.101 +/- 0.060 | 0.109 +/- 0.056 | 0.236 +/- 0.124 |
|  | Total | 0.076 +/- 0.045 | 0.022 +/- 0.012 | 0.100 +/- 0.054 |
| Tajima's D | Europe | -0.24 | **-2.37** | **-1.86** |
|  | YRI | -0.97 | **-1.73** | 0.78 |
|  | Total | **-1.52** | **-2.70** | -1.16 |
| Fu's Fs | Europe | -1.05 | **-26.78** | -4.84 |
|  | YRI | **-4.64** | -2.78 | 2.11 |
|  | Total | **-13.80** | **-25.17** | -4.76 |

In bold, significant values (p-value<0.05); total: all populations included; Europe: European populations only (Palestinians excluded).

The three PHAXs each show evidence of expansions across Europe starting around 20 KYA and pre-dating the Neolithic transition (beginning 10 KYA); in this, they are more consistent with the maternally-inherited mtDNA, which shows expansion ~15-20 KYA^9,10^ than with the male-specific Y chromosome, which shows expansions within the last 5 KY^9-11^. In turn, this behaviour is compatible with the female bias of X-chromosomal inheritance, and underscores the importance of male-specific behaviours in the recent reshaping of the genetic landscape in Europe.

Each of the small number of PHAXs we have studied here is independently inherited, yet together they present a reasonably consistent picture of prehistory. The X chromosome contains a large number of additional PHAXs (180 additional PHAXs identified across the whole X chromosome, excluding pseudoautosomal regions), and resequencing more would be desirable. In a much larger sample of PHAXs we might expect to find a broader distribution of behaviours, including possible recent expansion haplotypes reflecting the Bronze Age migrations. Statistical approaches designed for multilocus data, incorporating sequence data from ancient DNA, would maximise the insights these loci can provide about the past, and in particular that of females.

## Acknowledgements

We thank all DNA donors, and the NUCLEUS Genomics Facility of the University of Leicester for help with library preparation. This research used the ALICE High Performance Computing Facility at the University of Leicester. PMD was supported by a College of Medicine, Biological Sciences & Psychology studentship from the University of Leicester. CB, PH, DZ and MAJ were supported by a Wellcome Trust Senior Fellowship grant, number 087576, and MAJ and SB by Wellcome Trust Grant 057559.

Supplementary information is available on European Journal of Human Genetics’ website

